# Alternative approaches to single-cell trajectory inference using a commute time matrix

**DOI:** 10.1101/2025.02.12.635984

**Authors:** Douglas W. Houston

**Affiliations:** Department of Biology, The University of Iowa, 257 BB, Iowa City IA, 52242, USA

**Keywords:** Cell fate specification, Cell trajectory inference, Single-cell data imputation, Pseudotemporal ordering, Cell fate probability modeling, Commute time, RNA Velocity

## Abstract

Single-cell technology has enhanced the high-resolution analysis of dynamic developmental cell fate decisions. A number of elegant mathematical and computational approaches have been developed for using single-cell genomics data to identify gene regulatory events and cell state changes during embryonic development and related processes. These approaches are typically used in combination to model dynamic cell differentiation trajectories but have different underlying mathematical foundations. The extent to which commonly used algorithms for trajectory modeling, such as data imputation, pseudotemporal ordering, and cell fate probability modeling, might be derived from the same underlying approach has not been widely explored. This work describes the use of a matrix based on the commute time of a graph as a single consistent kernel for cell fate trajectory modeling. The commute time kernel is derived from significant eigenvectors of the pseudo-inverse of the graph Laplacian in a manner that preserves commute time. This kernel matrix is used directly in trajectory inference methods and recapitulates the results obtained using different algorithms using three benchmark datasets. Additionally, a comparison of commute time kernels between spliced and unspliced counts was effective for identifying populations of circadian-cycling progenitor cells in differentiating pancreatic endocrine cells. Overall, this work identifies the commute time kernel as a potential parsimonious measure for multiple aspects of trajectory inference.

## Introduction

Single-cell genomic profiling plays a critical role in characterizing cell state transitions and exploring gene regulatory interactions during dynamic developmental processes such as embryonic cell differentiation, tissue homeostasis, regeneration, cellular reprogramming, and disease progression. Researchers have developed a comprehensive framework, drawing inspiration from fields such as statistical physics and machine learning, to model and reconstruct the dynamic changes in transcriptional and other states as cells differentiate using single-cell genomics data [1–3]. These changes are commonly referred to as cell differentiation trajectories (following the textbook idea of a *state space trajectory*). For these analyses, single cells are statically sampled and, in most practical cases, a fraction of the genes expressed are successfully barcoded and sequenced. The exploration and visualization of data therefore often benefit from the imputation of ‘missing’ gene expression [4,5], although the imputed data is typically not considered appropriate for subsequent statistical analyses. Moreover, temporal and other relationships between cells are lost unless supplemented with lineage tracing or other cell labeling. However, these cellular relationships can be modeled to some extent by deriving a pseudotemporal ordering (distance) relative to a chosen initial state [6]. Additionally, modeling of RNA splicing dynamics, so-called ‘RNA velocity’ [7,8], is purported to add information to help orient the order of differentiating cells. Algorithms have also been developed to model random walk probabilities for cells adopting specific fates [9,10]. Combining these various computational approaches allows researchers to explore diverse cell fate trajectories and identify key gene expression trends.

Trajectory inference integrates the concept of ‘manifold learning,’ which involves creating an abstract representation of data as a lower-dimensional shape within the higher-dimensional ‘gene expression space’ [2]. Numerous interconnected algorithms, each with independent mathematical derivations and computational implementations, have been devised for manifold learning and applied to different aspects of single-cell trajectory inference [11]. Notably within these frameworks, however, algorithms are not usually employed interchangeably — imputation methods are not normally used for pseudotemporal ordering, and vice versa, and probability calculations are done using related but distinct methods. Navigating these different implementations and mathematical foundations can pose a challenge for new students in single-cell biology and those who may be lacking in-depth mathematical and computing backgrounds.

In addressing these challenges, the approach taken here involves deconstructing the main ideas of trajectory inference and reformulating them from a new direction. The goal is to better identify practical limitations and to learn first-hand the connections and commonalities among existing approaches. This work introduces a single graph-based reinterpretation of imputation, pseudotime, and probability measures, relying on a version of graph principal component analysis (PCA) of the commute time [12]. Commute time in this context represents the expected time for a random walk to traverse from one graph vertex to another and back [13–15]. This measure is analogous to the effective resistance (resistance distance) metric employed in electrical network analysis, both of which can be derived from the pseudo-inverse of the graph Laplacian/Kirchhoff matrix [16,17]. Commute time/effective resistance naturally represents a sum of random walks and has proven effective in various machine learning applications, including collaborative filtering, link prediction, graph sparsification, segmentation, and computer vision [18,19]. Whereas its use as a distance measure in clustering has been criticized [20], commute time has been suggested for pseudotemporal ordering [21], but is not widely implemented in single-cell genomics.

Critically for the context of cell fate trajectory analyses, the commute time kernel bears a resemblance to widely used Markov matrix-derived diffusion maps [12,22]. Markov transition matrices and Laplacian matrices share eigenvectors and have eigenvalues differing by a value of one. Also, decomposition of these matrices and scaling by eigenvalues can form a graph embedding, which for the pseudo-inverse Laplacian, implements a kernel PCA that preserves commute times [12,23]. This embedding encodes the vertices and topology of a corresponding hyperacute simplex, through graph-simplex bijection [24,25], implying that other features beyond just commute times may be also be conserved and represented. The commute time kernel thus incorporates various aspects of scale-free random walks and diffusion maps [21,22]. As a result, it has the potential to encode different features relevant to the analysis of cell fate trajectories.

The present work describes an empirically based commute time-derived approach to trajectory inference and is used in comparison to results from published and well-characterized datasets: epithelial-mesenchymal (EMT) transition in human cells [26], human bone marrow differentiation [9], and mouse pancreatic cell differentiation [27,28]. The (square rooted) commute time kernel produces results comparable to Markov affinity-based graph imputation of cells (MAGIC)-based imputation in addition to favorable comparisons to pseudotemporal ordering obtained by the diffusion pseudotime (DPT) and Palantir algorithms [9,26,29].

Moreover, estimation of cell fate probabilities using this approach aligns well with established methods. The combined implementation of these techniques not only reproduces established gene expression trends but also identifies potential novel candidate genes. Additionally, a nearest neighbor graph based on the commute time embedding offers a simple and intuitive method to visualize cell fate trajectories. Furthermore, when applied separately to spliced and unspliced counts, these methods suggest an alternate approach to RNA velocity, adding an additional tool in the analysis of cell differentiation dynamics.

## Results

### A commute time kernel for trajectory inference

The formulation of the commute time kernel resembles the construction of matrices used for critical aspects of trajectory inference, specifically sharing features with: a) the exponentiated Markov matrix of the MAGIC algorithm [26], used for data imputation, b) the formulation of diffusion pseudotime (DPT; [29]), used for pseudotemporal ordering, and c) the Markov fundamental matrix for absorbing random walks, used for probability estimations, as in Population Balance Analysis (PBA) and Palantir algorithms [9,10](see Methods). These commonalities suggested that a commute time-based measure could serve as a single, alternative approach encompassing multiple aspects of trajectory inference.

The commute time kernel is derived from an undirected, unweighted *k*-nearest-neighbor graph (*k*-NN graph) constructed from principal components, followed by the computation of the pseudo-inverse of the combinatorial/discrete Laplacian/Kirchhoff matrix of the graph (**L**^+^; **Fig. 1A**). The eigendecomposition of this matrix is performed, and the commute time embedding is then constructed by multiplying the square root of the pseudo-inverse eigenvalues by the corresponding eigenvectors. This operation, restricted to significant eigenvalues, effectively performs PCA on the graph [12]. Whereas the computational complexity of calculating the pseudo-inverse of the graph Laplacian can be challenging, an alternate approach to commute time involves using the reciprocal of the square root of the Laplacian eigenvalues. This approach is viable because the non-trivial eigencomponents of the Laplacian and its pseudo-inverse are identical, and their eigenvalues are inversely related, simplifying the computation process by performing the less complicated Laplacian eigendecomposition instead of performing a pseudo-inversion.

**Fig. 1.**
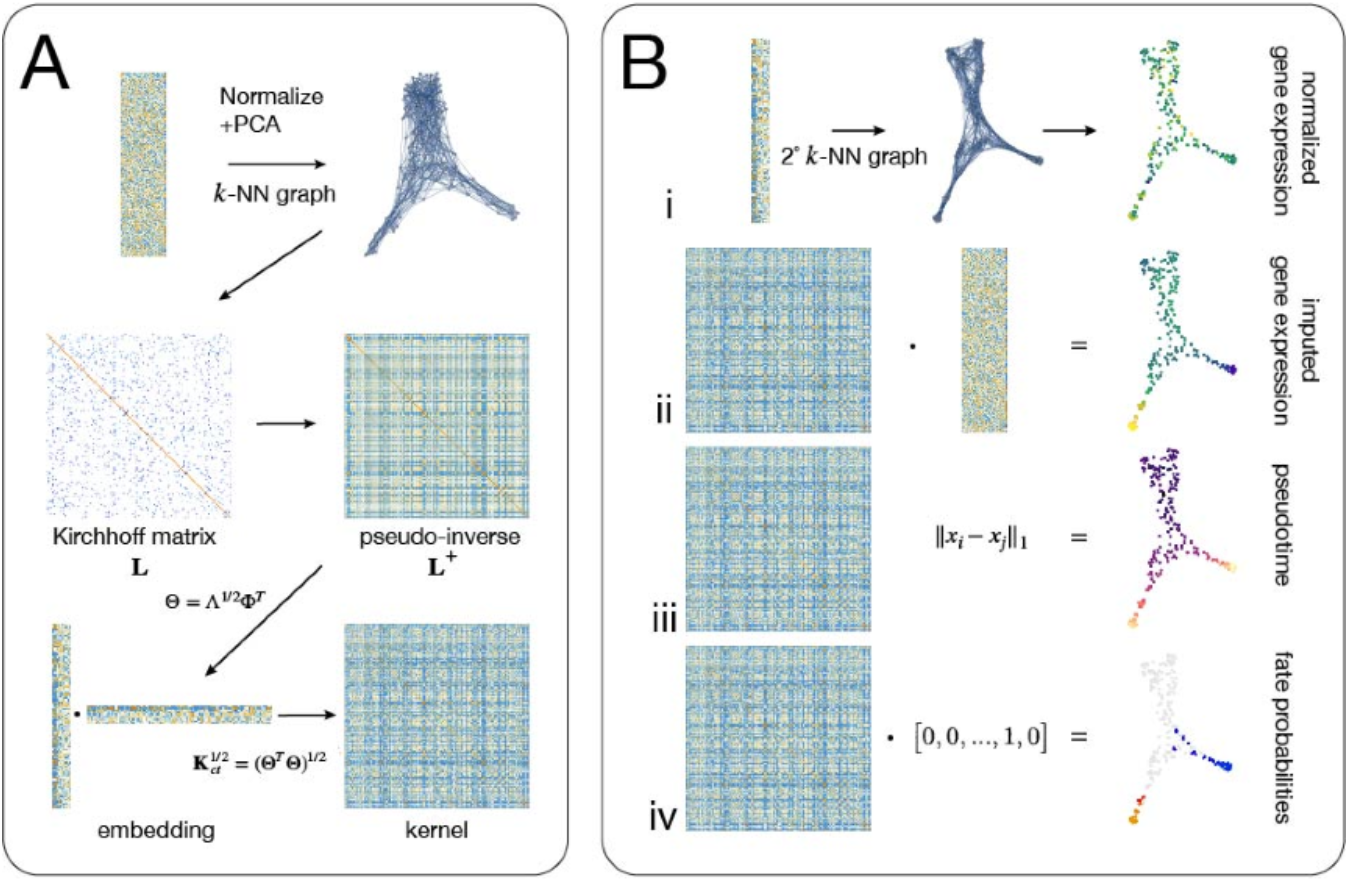
Graphical representation of the commute time method. (A) Data processing from a cells genes count matrix to a primary *k*-NN graph, which is decomposed via the pseudo-inverse of the graph Kirchhoff matrix to the matrix square root of the commute time kernel. (B) Applications of the commute time kernel. (i) The commute time embedding can be used to construct a secondary *k*-NN graph for visualization. (ii) Matrix multiplication of the kernel with the normalized data matrix can be used for data imputation. (iii) The L1-norm between vertices can be used to define pseudotime (iv) Matrix multiplication with unit vectors can define random walk probabilities for absorbing (differentiated) cell states. Data used for this example are from a simulated bifurcating differentiation process [10], representing 200 cells and expression of 50 genes.

The final commute time kernel is obtained by calculating the inner products of the vectors in the embedding matrix, creating a Gram matrix kernel. Subsequently, the matrix square root is applied (**Fig. 1A**). The embedding matrix serves an additional purpose, forming the basis for a secondary *k*-NN graph that can be used as an alternate way to visualize gene expression data and for other aspects of the data analysis (**Fig. 1Bi**). Imputation is applied through matrix multiplication of the commute time kernel matrix with the normalized count data, analogous to the procedure outlined in the MAGIC algorithm (Ref. [26]; **Fig. 1Bii**). Pseudotime is determined by computing the L1-norm of all cells with respect to the vertex corresponding to a designated root cell (**Fig. 1Biii**). Probability calculations involve multiplying the commute time kernel by unit vectors representing vertices defined as terminal (absorbing) states (**Fig. 1Biv**).

This commute time methodology shares similarities with existing methods, and arrives at similar measures by a different route, retaining many of the advantages, including robustness to noise, sampling density, and factoring in all ‘diffusion components’. Importantly, it can achieve these properties with less parameter tuning, such as estimates for Gaussian kernels and powering the diffusion operator, and providing a consistent matrix that can be applied across different analyses.

### Commute time kernel imputation approximates MAGIC data diffusion

To first evaluate the applicability of commute time kernel-based metrics for single-cell data imputation, the algorithm was compared to the analysis of epithelial-to-mesenchymal transition (EMT) conducted in the seminal MAGIC algorithm publication [26]. These data were derived from single-cell sequencing of transformed mammary epithelial cells treated with TGF-beta to induce EMT. Prior MAGIC-based imputation accurately captured changes in gene expression during EMT, notably including the inverse relationship between *CDH1* and *VIM* or *FN1* expression (*E-cadherin, Vimentin, Fibronectin*), the increase in levels of transcription factor *ZEB1* coincidently with these genes, and a transient increase in *MYC* [26,30].

Following data processing and graph generation, commute time-based imputation was initially attempted by directly multiplying either the pseudo-inverse of the graph Laplacian (**L**^+^) or the commute time kernel matrix (**K**_*ct*_), without any modifications, to the normalized EMT data expression matrix. This procedure resulted in elevated but flat gene interaction patterns when plotting the expression of EMT marker *VIM* against that of *MYC*, indicating a potential issue of over-smoothing (**Fig. 2A**). Because these matrices represent the summation of random walks, analogous to a large exponent number in the MAGIC algorithm, a measure was implemented to reduce the extent of imputation, namely taking the matrix square roots of these matrices. Using the square root of the pseudo-inverse Laplacian ((**L**^+^) ^½^) proved less effective than the original pseudo-inverse (**Fig. 2A**). However, applying the matrix square root to the commute time kernel 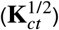 produced an imputation pattern for *MYC* plotted against *VIM* expression that closely resembled the results obtained using MAGIC imputation, showing a peak in *MYC* midway through EMT (**Fig. 2A**). Additionally, other gene expression relationships were captured in a manner similar to MAGIC (**Fig. 2B**). Analysis of the imputed values of eight representative genes (*MYC, VIM, FN1, EZH2, SNAI1, CDH1, CDH2*) showed a strong correlation, using Pearson Rho and Kendall Tau (rank correlation; Rho (*ρ*), Tau (*τ*)) measures, between data derived from MAGIC and the commute time method (mean *ρ*=0.97, mean *τ*=0.83), indicating a robust agreement between the two methods.

**Fig. 2.**
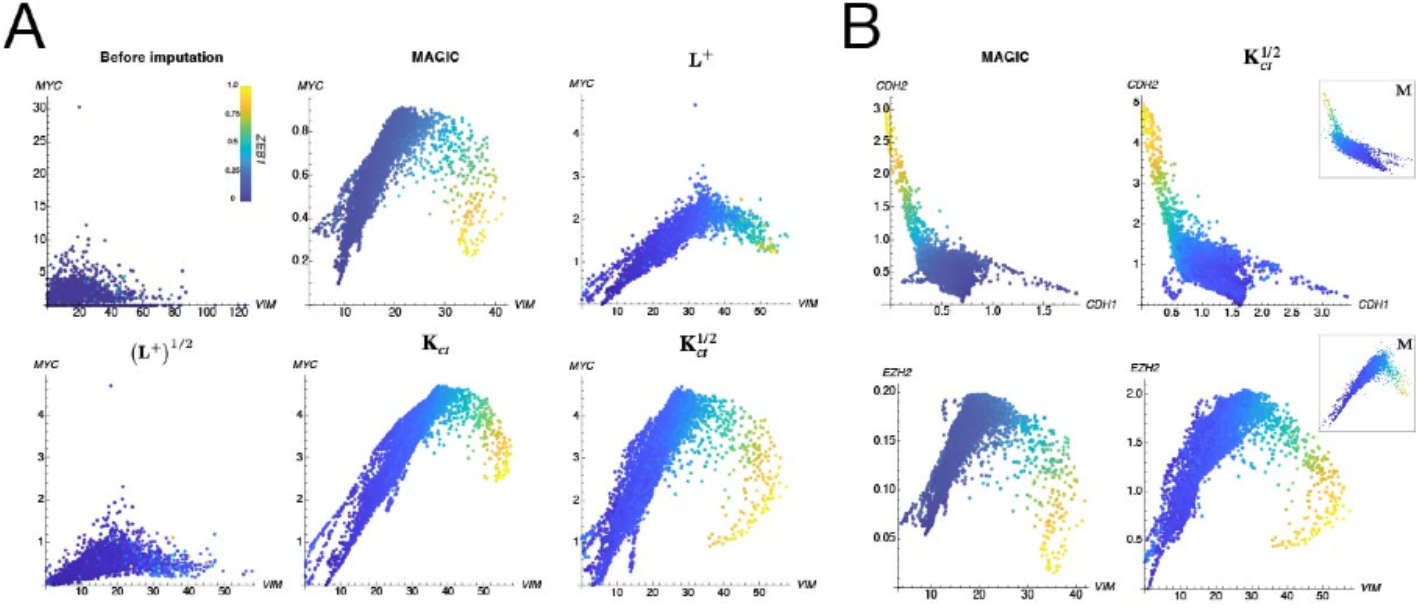
Commute time kernel-based imputation of gene interactions during EMT. (A) Scatterplots of MYC expression relative to VIM in single-cell sequencing of EMT. Normalized counts are shown before (Before imputation) and after application of different imputation operators. Cells are colored by the level of ZEB1 (as in [26], scaled), a gene important for EMT. VIM is a marker of the mesenchymal state. MAGIC (Markov affinity-based graph imputation of cells; [26]); **L**^+^, pseudo-inverse of Laplacian matrix; **K**_*ct*_, commute time kernel. Power of ½ is the matrix square root. (B) Additional comparisons of commute time with MAGIC using different gene interactions during EMT. Insets show a comparison to imputation using the Diffusion Pseudotime (DPT) **M** matrix [29].

To explore the generality of this approach, an alternate computation of summed random walks was also tested. The matrix devised for diffusion pseudotime (DPT) is a Markov transition matrix, with the contribution of the constant eigenvector subtracted [29]. This operation transforms the matrix into a scale-free form, **M**, representing the sum of all random walk lengths. This approach is thus comparable to commute time, which by construction, lacks the contribution of the constant eigenvector, and also represents a sum of random walks. To test this alternate derivation, the row normalized, symmetrical transition matrix was generated from the EMT graph and the dominant eigenvector was subtracted, following the procedure for DPT [29]. The resulting matrix **M** was then directly used for imputation. The results were visibly similar to those obtained with the commute time kernel and demonstrated good correlation using the same gene sets (insets in **Fig. 1B;** mean *ρ*=0.92, mean *τ*=0.71). This concordance using an alternate scale-free matrix further supports the efficacy of this approach for imputation in the context of single-cell data analysis.

The quantification of gene interactions (or lack thereof) in the context of EMT progression and the potential for regulation by ZEB1 were successfully recapitulated using the commute time methodology. These interactions could be similarly quantified using a measure of mutual information based on conditional probability (DREMI; [26,31]; **Fig. S1**). The commute time method was also robust to 95% artificial dropout and 80% down-sampling, similar to the results obtained with MAGIC (**Fig. S2**) [26].

Overall, these results suggest that effective imputation of single-cell sequencing data can be achieved through the de-exponentiation (square rooting) of the commute time kernel, or through other scale-free affinity matrices in general, analogous to the optimal exponentiation of a Markov transition matrix.

### Commute time kernel and pseudotemporal ordering

A second, more complete trajectory analysis using multiple measures derived from the commute time kernel was performed using a human bone marrow differentiation dataset, analyzed as reported ([9]; replicate number one). Imputation based on the bone marrow-derived commute time kernel successfully identified major cell lineages based on imputed gene expression, as shown in force-directed/spring-electrical embedding plots of the primary *k*-NN graph (**Fig. 3A, 3C, Fig. S3A**; e.g., *GATA1* in the erythroid lineage). Furthermore, this analysis revealed known gene expression patterns during cell fate bifurcations such as the erythroid vs. megakaryocytic, neutrophil vs. dendritic cell, and erythroid vs. myeloid cell differentiation (**Fig. S3B**; [9]).

**Fig. 3.**
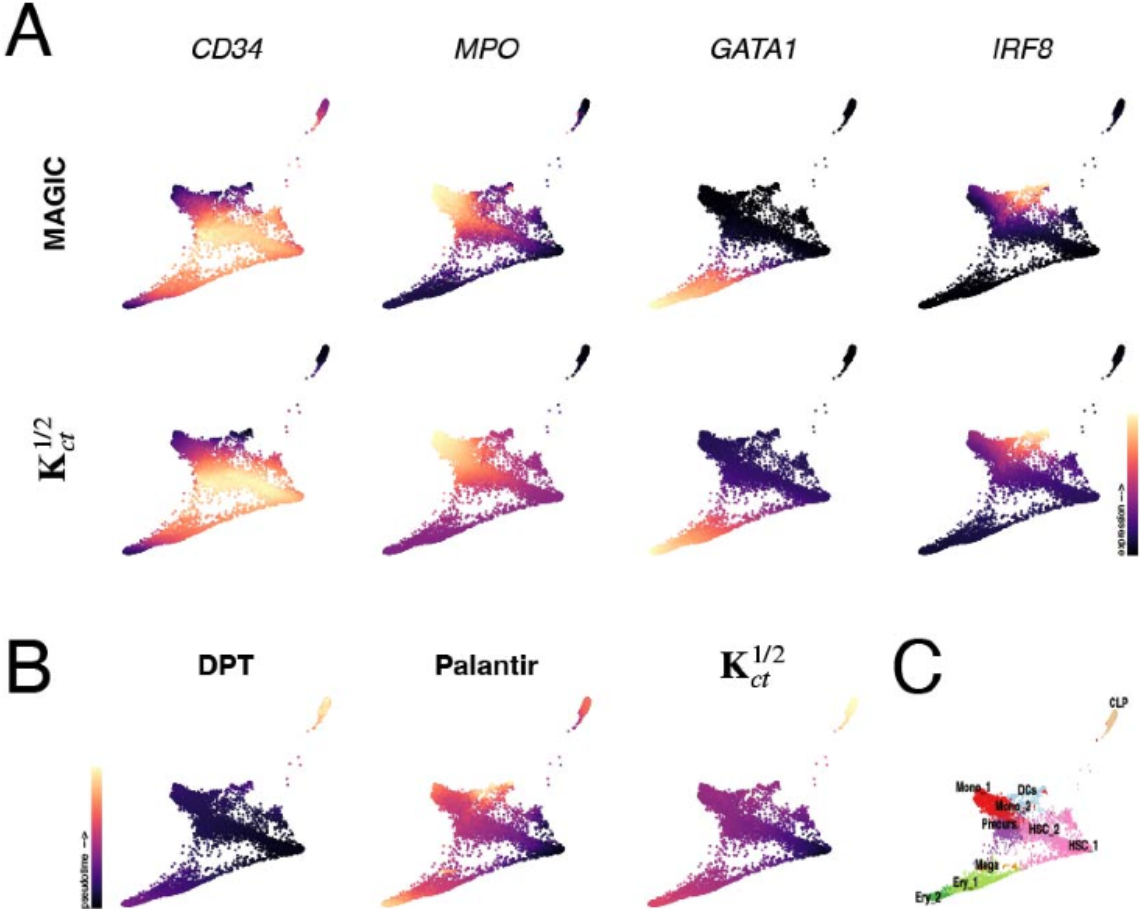
Commute time imputation and pseudotime applied to human bone marrow differentiation. (A) Force-directed graph plots of imputed expression for bone marrow stem cells/progenitors (*CD34*) and selected lineage-specific markers (myeloid, *MPO;* erythroid, *GATA1;* dendritic, *IRF8*). Cells are colored by imputed expression levels. Results using MAGIC and the commute time kernel 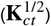 are shown. (B) Pseudotime comparison using DPT, Palantir and commute time, using the same root cell in the hematopoietic stem cell population. Cells are colored by scaled pseudotime (from 0-1). (C) Location of cell clusters on the graph, as determined by Setty et al. (Ref. [9]). These plots were generated using a modified Python workflow.

To facilitate trajectory analysis, a common pseudotime was defined using a commute time-based pseudotime compared to published DPT and Palantir values. DPT uses the scale-free matrix **M** (see previously, [29]), or an ‘on-the-fly’ equivalent derived from the diffusion map [21], to compute Euclidean distances between vertices and a defined root (starting) cell; longer distances thus representing longer time. Palantir is based on a different approach, involving shortest path random walks on a diffusion map-based *k*-NN graph [9].

Initially, the pseudo-inverse Laplacian matrix **L**^+^of the graph, which can express the commute time, was (naively) tried directly in place of the DPT matrix to compute distances, using the same root cell. **L**^+^ produced a pseudotime reasonably correlated to both DPT (*ρ*=0.83; *τ* =0.79) and to Palantir (*ρ*=0.74, *τ*=0.72). However, these correlations were improved with the commute time kernel 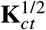 and, empirically, using the L1-norm (rather than the L2-norm) to increase robustness in high dimensions, producing visibly similar pseudotimes (**Fig. 3B**). These alterations did not alter the correlations with DPT but increased the association with Palantir (vs. DPT: *ρ*=0.81; *τ*=0.79; vs. Palantir: *ρ*=0.90; *τ*=0.76). Thus, the commute time kernel proves comparable to existing measures in formulating a rooted pseudotime.

Visualization using a commute time embedding-based *k*-NN graph offered a more intricate view of differentiation trajectories. This secondary graph showed a detailed continuous topological view of the relationships between cells during differentiation, characterized by extended arm branching reflecting the different differentiation states. Terminal states for each lineage manifest at the termini of the graph tips, as evident in the expression of terminal marker genes (e.g., *ITGA2B* in the megakaryocytic lineage; **Fig. 4**; imputed). The commute time-based graph embedding revealed several lineages with greater resolution, including terminal erythrocytes, subgroups of monocytes, and lymphocyte precursors (**Fig. 4**).

**Fig. 4.**
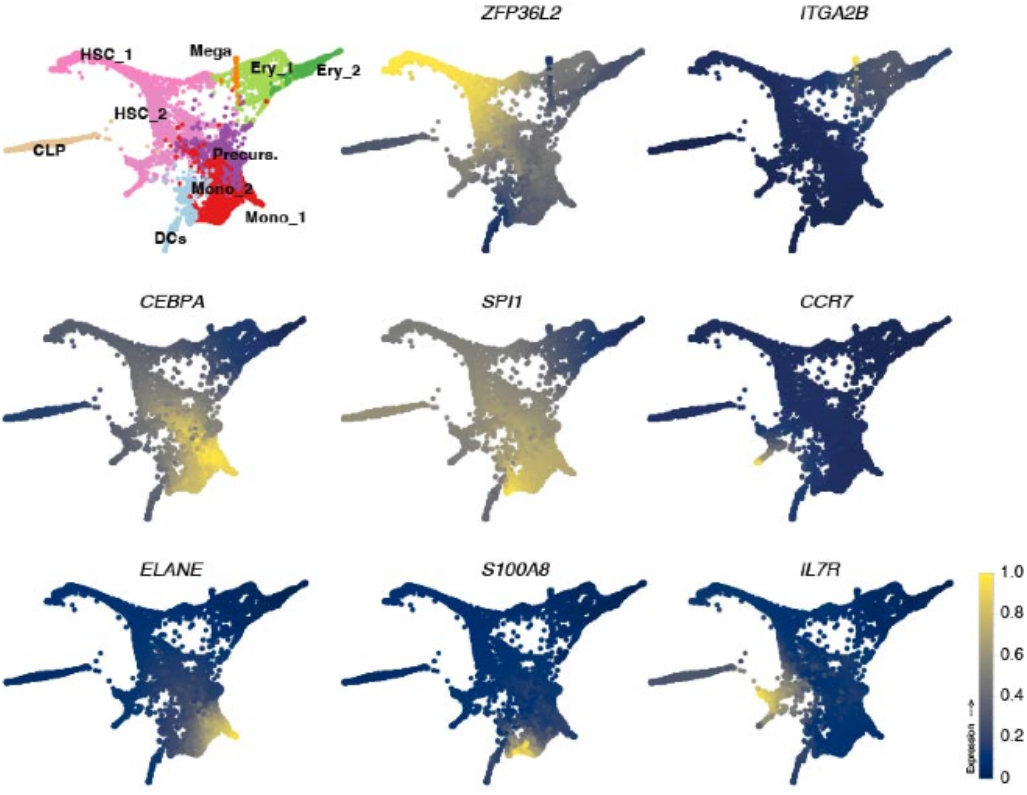
Visualization of single-cell data using commute time embedding-derived nearest neighbor graphs. Graph plots (spring-electrical layout) derived from the commute time embedding of bone marrow differentiation single cell data. The clusters from Fig. 3C are highlighted in the upper left panel. The remaining panels show imputed expression (scaled) for stem cell (*ZFP36L2)* and various lineage marker genes.

An important aspect of differentiation trajectory inference involves probabilistic modeling of cell fates using random walk probabilities with absorbing states (i.e., the differentiated states). To adapt this process using commute time, terminal states were selected manually, using the SPRING browser as in the PBA method [10,32]. Probabilities were estimated from the square rooted commute time kernel using a modified, simplified, escape probability calculation [33,34]. In this analysis, probabilities gradually increase along the graph arms, with higher probabilities assigned to the extreme tips (the selected cells) of the graph arms **(Fig. 5A)**, representing the most differentiated cells. The commute time graph exhibited several additional fates, but in the seven cases aligning with those identified by Palantir the probability values showed robust correlation with those Palantir algorithm probabilities (**Fig. 5A**). These probabilities were used to assign weights to non-linear models of lineage-enriched gene expression across pseudotime to delineate gene expression trends [9]. Trends observed for markers *GATA1* and *SPI1* using commute time measures closely mirrored those computed from published MAGIC-imputed data used with Palantir derived probabilities **(Fig. 5D)**. Conditional probability density plots of these genes (see previously) additionally revealed the corresponding trends in their expression patterns over pseudotime (**Fig. 5E**). Thus, the commute time matrix can be employed effectively across multiple aspects of trajectory inference.

**Fig. 5.**
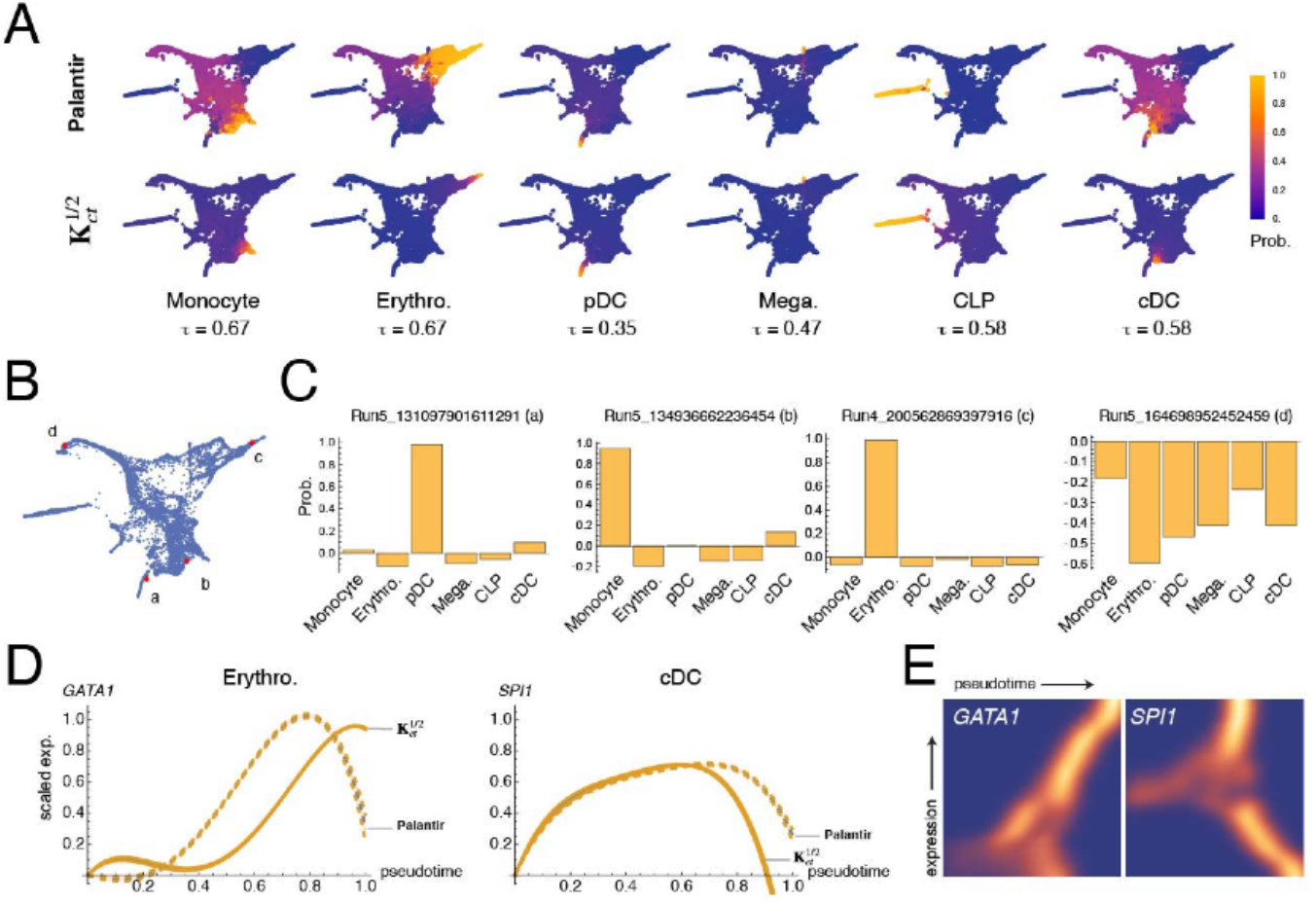
Commute time usage in cell fate probability and gene expression trend analysis. (A) Force-directed graph plots of modeled cell fate probabilities obtained using Palantir and commute time. Kendall’s tau values (*τ*, rank correlation coefficients) are shown below each pair of plots. Kendall Tau Tests for significance resulted in p=0.0 in each case. (B) Selected cells for individual probability assessment (taken from [9]). (C) Fate probabilities for cells in the (a) pDC dendritic lineage (plasmacytoid dendritic cells); (b) monocyte; (c) erythroid; and (d) stem cell lineages. (D) Gene expression trends for *GATA1* (weighted by erythroid probability) and *SPI1* (weighted by cDC (conventional dendritic cell) fate). Trends were calculated using imputed expression and pseudotime derived from commute time (solid lines) and from MAGIC imputation/Palantir pseudotime (dashed lines). (E) Density plots of conditional probability for gene expression across pseudotime; *GATA1* shows higher probability of high expression late in pseudotime (left panel), whereas *SPI1* (right panel) bifurcates into high and low-expressing lineages around 0.6 normalized pseudotime (the former corresponding to the cDC lineage).

### Commute time kernel analysis reveals novel candidate genes in pancreatic islet cell differentiation

In a third illustrative example, the commute time algorithm was used to analyze a dataset of E15.5 mouse pancreas development [27,28,35]. Additionally, the method was extended by using the respective commute time kernels to analyze data from spliced versus unspliced RNAs (see subsequently). Visualizing the total count data using the secondary *k*-NN graph revealed a smooth and tightly represented graph showing distinct resolution of all relevant pancreatic lineages (**Fig. S4**). Ductal cells and early progenitors expressing *Sox9* and *Neurod2*, respectively, were positioned along one extended arm of the plot, whereas *Fev*-expressing transition cells occupied the central region. Additional radiating arms representing the various differentiated endocrine cell fates were also evident (**Fig. S4**).

Importantly, delta and epsilon islet cell fates became well-resolved using this secondary commute time graph, as indicated by enrichment of *Sst* and *Ghrl*, respectively. In addition, Pancreatic Polypeptide (PP)/gamma cells, which express the peptide hormones *Pyy* and *Ppy* (**Fig. S4**) could be distinctly identified. Notably, a prior analysis of this dataset using an RNA velocity-based approach (CellRank, Ref. [28]) did not delineate the delta cells as well and failed to specifically identify the gamma-lineage, which overlapped the *Fev-*expressing cells and delta cells in that study. To assess the effectiveness of the commute time approach for identifying critical delta and gamma lineage genes, gene expression was correlated with trajectory probabilities. Included in the top 50 highest correlated genes were known delta cell lineage markers, *Sst, Rbp4, Hhex* and *Cd24a*. Also, novel delta-related genes found by CellRank were also identified, such as *Hadh* and *Arg1* [28]. Other new delta-enriched genes indicated by the commute time trajectory included *Fos* and *Junb, Ptprz1, Igfbp5*, and *Mest* (**Fig. 6A**). These genes have not been explicitly studied in developing delta cells but have been implicated in pancreatic function and in pancreatic cancer (e.g., [36–38]).

**Fig. 6.**
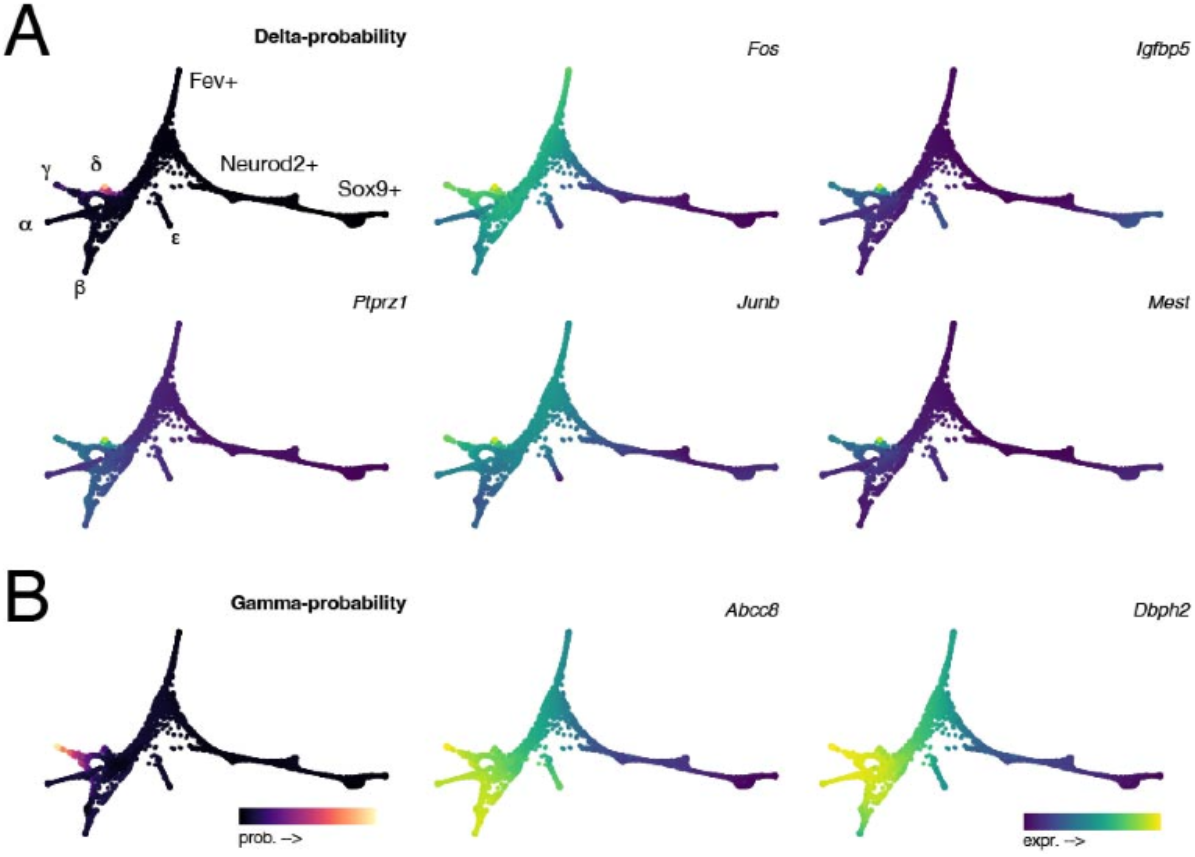
Novel delta and gamma lineage marker gene expression. Force-directed graph plots of E15.5 pancreatic cells showing predicted probabilities for (A, upper left) delta lineage fate and for (B, left) gamma lineage fate. (A) Expression of genes with high correlation with delta fate. (B) Expression of genes with high correlation with gamma fate. Positions of different endocrine lineages and progenitors are labeled in (A).

Fewer unique gamma islet-expressed genes could be identified with high confidence, owing to gamma cell heterogeneity. *Pyy*, a canonical gamma lineage marker, was most the correlated gene with gamma lineage probability and was also the most significantly enriched gene in gamma cells using differential gene expression analysis [55]. However, manual inspection additionally identified enrichment of two candidate genes from the correlation list that are implicated in regulating gamma and alternate islet cell fates and function: *Abcc8* and *Dbpht2* (**Fig. 6B**). Gamma cells are thought to contribute to the beta cell population. Deletion of *Abcc8* in adult beta cells led to those cells switching fate to that of gamma/PP cells [39], suggesting this *Abcc8*+ population might represent those that would later contribute to the beta cell pool. Similarly, *Dbpht2* has been shown to be enriched in an early differentiating beta cell population [40]. *Dbpht2* is also expressed in alpha cells (**Fig. 6B**) and may also have a role in their differentiation. Overall, these analyses show the efficacy of using the commute time kernel approach in characterizing cell differentiation dynamics and in providing new opportunities for gene discovery.

### RNA velocity-like analysis of islet cell differentiation using between graph pseudotime

Last, to gain additional insight into differentiation dynamics from comparing spliced versus unspliced transcripts, the data for each transcript set in the pancreas data was processed individually and commute time kernels and graphs were generated for each. The intuition behind this approach is that if the splicing dynamics are similar across all cells, then the spliced and unspliced graphs should be similar to the overall graph (and to each other). Alternatively, cells with unique sets of unspliced gene expression, i.e., those in early stages of differentiation, would be expected to contribute to a different graph structure.

Processing of the pancreas data using spliced transcripts produces a graph similar to that of the overall transcripts, as expected (**Fig. 7A-C**, compare to **Fig. S4**). However, the unspliced graph is different, with one prominent arm extending from a more condensed mass of cells (**Fig. 7D-F**). Cells in this arm express beta cell markers (e.g., *Insulin2/Ins2*; **Fig. 7A, D**), suggesting a relatively greater extent of ongoing new gene transcription in this lineage. Other cell types occupied distinct territories in the main body of the graph but were not resolved in fine detail (e.g., *Gcg*; **Fig. 7B, E**). To represent the relative difference in the contribution of unspliced RNAs, a method analogous to pseudotime *between* graphs was computed. This method uses the commute time kernels from the two graphs to find the distance between corresponding vertices (splicing pseudotime). The beta cell lineage exhibited the greatest distances between spliced and unspliced vectors corresponding to the same cells (**Fig. 7C, F**), reflecting the visible differences in the two graphs.

**Fig. 7.**
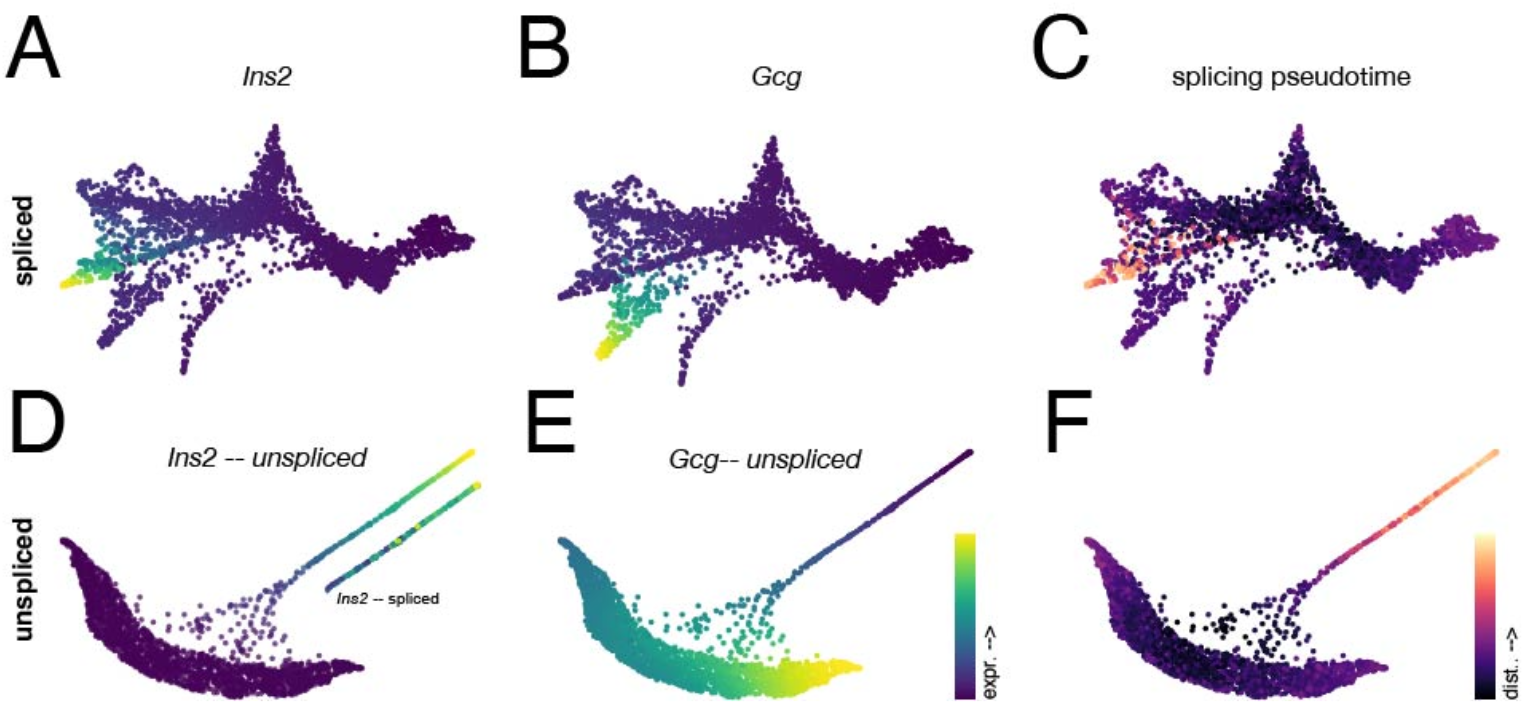
Cell state trajectory analysis using distance between nearest neighbor graphs of spliced and unspliced transcripts. Force-directed graph plots of E15.5 pancreatic cells derived from individual *k*-NN graphs of (A-C) spliced and (D-F) unspliced transcripts. (A, D) Imputed expression of spliced and unspliced *Ins2* expression, plotted on the respective graphs. (B, E) Imputed expression of spliced and unspliced *Gcg* expression, plotted on the respective graphs. (C, F) Visualization of splicing pseudotime, the scaled L1-Norm between corresponding row vectors of the commute time kernels for the respective spliced and unspliced datasets. Cells with the greatest distances are in the beta cell lineage, and are localized to the tip of the extended beta cell arm in the unspliced graph.

To identify actively transcribed genes contributing to this difference, normalized *unspliced* counts were correlated with this splicing pseudotime measurement. In addition to known beta cell genes *Ins2, Nnat* and *Iapp*, other genes related to beta cell maturation and insulin secretion (e.g., *Ppp1r1a, Gng12, G6pc2, Slc2a2;* **Fig. 8A**, top) were also among the top correlated genes differing in unspliced vs. spliced transcripts. Additionally, genes implicated in controlling circadian rhythm were well-correlated with splicing pseudotime (e.g., *Phactr1, Rora;* [41,42], **Fig. 8A**, bottom). When visualizing unspliced counts (imputed) on the graph, beta cell maturation genes tended to be expressed at the tips of the graph arm (see representative examples in **Fig. 8A**, top, insets), whereas core circadian rhythm genes (*Clock, Per2, Cry2*; [43]) tended to be more highly expressed mid-way along the arm (**Fig. 8A**, bottom, insets).

**Fig. 8.**
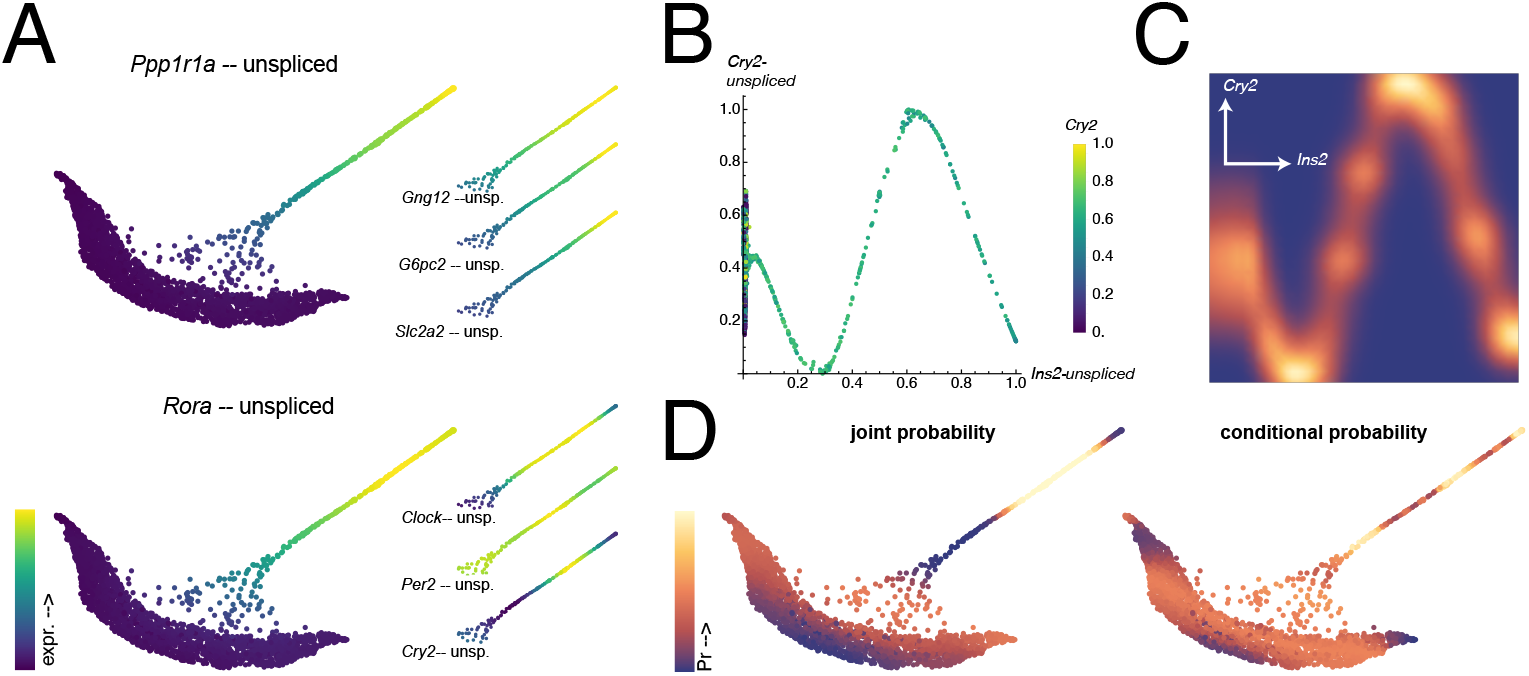
Enrichment of unspliced transcripts related to insulin secretion and circadian rhythm in differentiating beta cells. (A) Force-directed graph plots based on unspliced transcripts showing unspliced expression of genes involved in insulin secretion (upper panel: *Ppp1r1a; Gng12, G6pc2, Slc2a2* in insets) and circadian rhythm (lower panel: *Rora; Clock, Per2, Cry2* in insets) in the beta cell lineage. (B) Scatterplots of relationship between unspliced *Ins2* and *Cry2* expression (scaled expression; unscaled *Cry2* << *Ins2*). Cells are colored by spliced *Cry2* expression, which is at intermediate levels at all *Ins2* levels. (C) Density plot of conditional probability of *Cry2* levels on *Ins2* levels. (D) Unspliced graph plots showing joint probability (left, co-occurrence of *Cry2* and *Ins2*) and (right, conditional *Cry2* probability). As in (C), cyclical high probability at increasing unspliced *Ins2* levels is evident.

Mature adult beta cells are known to undergo circadian oscillations in Insulin secretory activity [52, 53]. Also, analysis of *Ins2::gfp* knockin mice showed that cells with high *Ins2* transcription are enriched in both beta cell maturation and circadian clock related transcripts, suggesting that beta cell heterogeneity at least partially results from temporal dynamics [54]. The cells found in the main prominent arm of the unspliced graph could thus represent this population and suggest this pathway is active embryonically, at the onset of beta cell differentiation. Circadian rhythm genes showed cyclic expression when plotted against unspliced *Ins2* transcripts (e.g., *Cry2;* **Fig. 8B**) and could alternatively be visualized and quantified using the DREMI-based approach (see previously, and [26,31], **Fig. 8C**). Additionally, plotting the joint and conditional probabilities of unspliced *Ins2* and *Cry2* identified regions of overlap (**Fig. 8D**), which could indicate key points of circadian regulation.

These analyses thus suggest alternate methods for using unspliced transcript data in single cell experiments that do not depend on modeling splicing dynamics and other constraints of RNA velocity, as typically implemented.

## Discussion

The primary insight from this study is that a single matrix derived from the pseudoinverse of the graph Laplacian can be used equivalently to three separate, interrelated, but similar single-cell data analysis measures: the exponentiated transition matrix of MAGIC imputation [26], the scale-free transition matrix of DPT [29], and the Markov fundamental matrix in absorbing probability (cell fate) calculations [9,10]. The aim of this work was mainly autodidactic, to explore the rationales behind single-cell analysis algorithms and re-express them in a different way to promote understanding. The math behind various single-cell analysis methods is represented in an accessible form using Mathematica notebooks to encourage experimentation and tinkering with parameters. Symbolic math languages are not widely used in bioinformatics but can useful as a bridge between abstract mathematical ideas in papers and opaque, pre-packaged computational pipelines. The workflow presented here is based on the various original derivations, for which this work is indebted. The commute time method would require additional mathematical formalizing to be more completely compared with previous methods. In the spirit of C.H. Waddington (1940, Ref. [44]), writing on his own mathematical dabbling, this paper will “…have fulfilled its function if its inelegance annoys some mathematician sufficiently [to improve upon it].”

The commute time kernel measure developed in this study has been widely used in other contexts but has not been previously reported in single-cell analysis. One possible conceptual finding here is that subinfinite random walks, using the commute time kernel matrix square root or the diffusion pseudotime (DPT), can form the basis for both imputation operators and pseudotime measures. The spectral similarity between Markov and Laplacian matrices positions the current method as analogous to a reduced Laplacian eigenmaps/diffusion maps method [22,45,46], although with different weighting of the eigenvectors in the construction of the Gram matrix kernel, representing the sums of random walks.

Commute time has been criticized as a distance metric, as it converges to the sum of vertex degrees for random graphs [15,20]. In real-world settings, commute time/resistance distance has proven useful [47,48], and discrepancies between commute time and vertex degree of up to 30% exist for non-random data [20], suggesting this convergence may not be a practical limitation. Regardless, the methods described here do not involve commute time directly, but operate on a commute time-preserving matrix. Square rooting on this matrix is more favorable in practice than using the non-square rooted commute time or pseudoinverse Laplacian directly and may be necessary to reduce the extent of random walk summation.

The visualization method using the *k*-NN graph representing simplex coordinates aligns conveniently with the commute time matrix used here. Whether this correspondence is coincidental or reflects deeper relationships between single-cell measures and simplex geometry warrants future investigation. Notably, the commute time kernel is not used for clustering in the current method, but its geometry could be effectively applied in simplex-based clustering algorithms, e.g. CellRank (Ref. [28]).

While the goal of this work wasn’t necessarily to devise a ‘better’ analytical framework, the commute time method does effectively recapitulate prior results while revealing some new information. In the bone marrow and pancreas datasets, the commute time embedding graph, using force-directed graph plotting, delineated lineages more clearly than the usual UMAP plots. Notably, commute time-based plotting identified known cell types that were poorly separated in other representations. In the bone marrow, these cell types included early lymphoid progenitors expressing *IL7R* or *CCR7*, as well as different monocyte populations. In the pancreas, the delineation of gamma and delta cell lineages facilitated the identification of novel genes in those lineages. This study used several previously well-characterized datasets. It will therefore be important to assess the efficacy of the commute time method in the context of novel experimental data, where the outcome is largely unknown.

This work also explores a novel use of spliced versus unspliced count data from single-cell RNA sequencing experiments. By generating graphs from each dataset, differences in transcript abundance between spliced and unspliced forms can be used to create distinct graphs, exemplified in the pancreas data in which beta islet cells feature prominently in the unspliced graph. Additionally, the differences in the graphs can be represented by a new ‘splicing pseudotime’ measure, analogous to the distance between the same cell in the spliced vs. the unspliced graph. This distance is expected to be greater in cells overrepresented in unspliced transcripts, which would be those undergoing the most rapid differentiation. This distance may thus represent a simpler alternative to the so-called RNA velocity, which is controversial for being highly sensitive to distortions based on the choice of embedding parameters.

It’s not clear how well the splicing pseudotime compares with RNA velocity, given the differing assumptions and variability with the latter, although both methods suggest strong indications of beta cell differentiation in the pancreas dataset. The splicing pseudotime method also identified the link between circadian rhythm genes and insulin secretion in early beta cells, which had been previously observed experimentally in adult pancreas but was not readily discerned in other single-cell analyses of embryonic islets. More examples would be needed to determine the full utility of splicing distance in identifying differentiating cell populations.

An almost limitless variety of increasingly complicated computational models could conceivably be applied to single-cell genomics data. However, the trend towards mathematical sophistication should be balanced with real-world applicability and general comprehensibility. As single-cell genomics evolves from large-scale cell-atlas projects to more routine use in many settings, experimental scientists will increasingly need a more intuitive grasp of the relevant mathematical ideas and computational data analysis techniques. Helping the next generation of scientists build this intuition is necessary, as future research in developmental biology will greatly depend on expertise in both the computational and experimental worlds, without necessarily seeing them as totally separate disciplines.

## Methods

### Data processing and graph construction

Raw count matrices (*C* = *mxn*; cells by genes) were imported and used without further filtering. Counts per cell were L1 normalized and scaled using the median count per cell, based on Dijk et al. [26]:

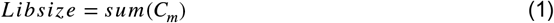

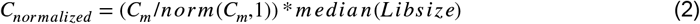

Log-transformation and selection of highly variable genes were not performed. Principal components analysis (PCA) was used as an initial linear dimension reduction, keeping the components representing ~70-85% of the total variance (number of pcs = 50, 50, and 30 in the examples used here). The results are robust to the exact numbers of PCs used. PCs can be determined using the covariance (zero centered data) or correlation (standardized data) matrices; the latter was used for the bone marrow differentiation analysis in this study.

An initial *k*-nearest neighbor (*k*-NN) graph is generated from the reduced data using a *k* of approximately the square root of the number of PCs. There is some flexibility around this choice; it is recommended to use a value for *k* resulting in a connected graph. This primary graph is then subjected to spectral analysis to derive the various metrics and can be used directly for visualization using spring-electrical embedding plot.

### Spectral decomposition and commute time kernel construction

Commute times/effective resistances are computed from the pseudo-inverse (Moore-Penrose inverse) of the graph Laplacian matrix (**L**^+^). Eigen-decomposition of **L**^+^ leads to a non-linear dimensional reduction and embedding that preserves commute times and, after retaining significant eigenvalues, is analogous to kernel PCA on this matrix [12]. The commute time kernel is the matrix of inner products (Gram matrix) of this embedding, the matrix square root of which is used here in trajectory analyses.

Briefly, the unnormalized/combinatorial Laplacian/Kirchhoff matrix **L** of the initial graph was computed as the degree matrix **D** (the diagonal matrix of graph vertex degrees *d*_*i*_) minus the adjacency matrix **A** (**L** =**D** − **A**) using a built-in Mathematica function (KirchhoffMatrix; [49]). The Moore-Penrose inverse/pseudo-inverse of this matrix can be readily determined for small datasets but can be computationally challenging for large matrices. However, because the eigenvectors are conserved between **L** and **L**^+^, the reciprocal of the Laplacian eigenvalues (*λ*^−1^) can be used equivalently. In small-scale experiments, directly using the pseudo-inverse was preferred since it is more intuitive to retain the highest eigencomponent pairs after the decomposition. The eigendecomposition of **L**^+^ can be represented by equation (Eq. 3) and its quadratic form (Eq. 4):

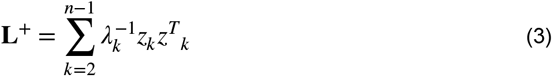

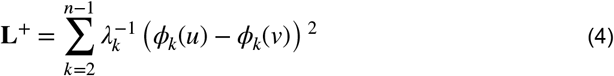

(Eq. 4) represents the commute time between vertices *u* and *ν*, or the expected (average) number of steps to visit vertex *ν* and return, beginning at vertex *u*. The pseudo-inverse eigensystem is well-known to comprise a commute time embedding, or mapping the data to a Hilbert subspace in a manner that preserves the commute times and represents a topological simplex of the graph. For the unnormalized/combinatorial Laplacian, this embedding becomes [12,15,22]:

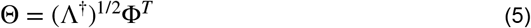

wherein Λ^†^ is the inverse of the diagonal matrix of the Laplacian eigenvalues (*λ*, sorted in increasing order) on the diagonal and **Φ** is the eigenvector matrix in standard representation. The Gram matrix kernel of the embedding, the commute time kernel **K**_*ct*_, is then obtained through the inner product of the embedding with itself, which restores the basic form of **L**^+^ (a Gram matrix and valid kernel in its own right), with the smallest eigenvalues discarded:

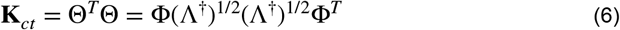

The final kernel is obtained by taking the matrix square root 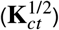, as was determined empirically, and the resultant matrix used for imputation, pseudotime, and probability determination.

### Visualization based on the commute time embedding

A secondary *k*-nearest neighbor (*k*-NN) graph was generated from the commute time embedding (Θ^*T*^) heuristically using a *k* of approximately one-half to two-thirds the square root of the number of cells.

### Commute time kernel-based data imputation

Imputation was applied through matrix multiplication of the commute time kernel with the normalized data matrix:

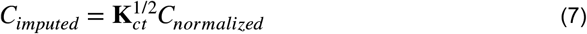

The imputed data matrix is then scaled as in the MAGIC algorithm (Eq. 8, Ref. [26]) to restore gene expression levels close to original levels, and then shifted to eliminate negative expression values.

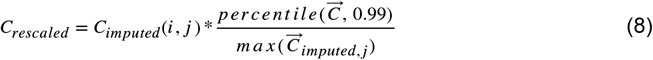

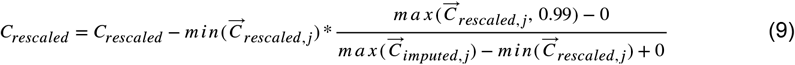

The commute time kernel recapitulates several aspects of MAGIC, including applying an optimal amount of diffusion to the data, decreasing the high-frequency (noisy) information by eliminating trivial eigenvalues, and providing non-linear weighting of close versus distant vertices through the commute time distances, but doing so by a different process.

For comparison to another scale-free measure, the **M** matrix of diffusion pseudotime (DPT) was generated as described [29]:

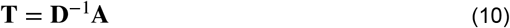

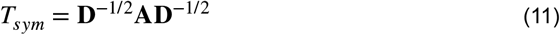

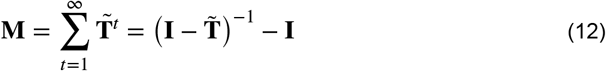

In (Eq. 12), **I** is the Identity matrix, and 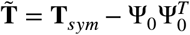, with **Ψ**_0_ referring to the leading eigenvector (i.e., the constant eigenvector corresponding to the largest eigenvalue of 1.0) of **T** (as in Ref. [29]). This matrix was used for imputation as in (Eq. 7).

### Pseudotemporal ordering

Because of the analogies between the commute time and diffusion maps, pseudotime was computed by calculating distances using the L1-norm from a preselected root cell (*i*) to all other cells using the kernel matrix:

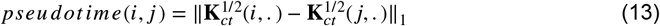

### Mutual information and conditional mutual information

The mutual information (Eq. 14) is defined as the reduction in uncertainty of a random variable (entropy) by knowing another random variable (conditional entropy) [31]:

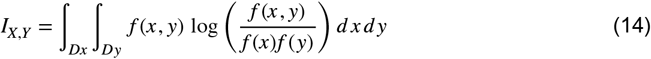

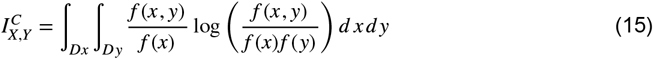

*f* (·) represents a probability density function. DREMI (Eq. 15) rescales the mutual information by the conditional probability density function to better identify relationships between variables [31].

### Probability estimation

Cell fate probabilities were estimated directly using the commute time kernel using an escape probability calculation modified from [50]. Escape probability is the probability of a random walk starting at vertex *w* and visiting *u* before ‘escaping’ to *ν*, with *ν* being taken as an absorbing state (i.e., a terminal cell state). Here, escape probability is used as escape to any other vertex and is computed by multiplying the commute time kernel matrix by a unit vector (**e**_*i*_) with a value of one at the absorbing state(s) and zeros elsewhere.

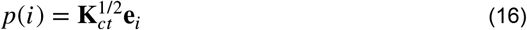

Several cells in each terminal state are analyzed and the escape probabilities averaged. Alternatively, the commute kernel can be multiplied by a matrix of unit vectors, one column for each absorbing state, an operation closely related to the usual probability calculation using the Markov fundamental matrix.

Terminal states were chosen manually by exporting graphs to SPRING [32] and selecting cells at the tips of the extended arms, but the commute time procedure would be compatible with other methods for finding terminal states.

### Spliced vs. unspliced transcript analysis

To reflect differences between graphs derived from spliced and unspliced counts, a splicing pesudotime was implemented based on the L1-norm distance between the same vertex vector (*i*) in commute time kernels derived from spliced (*s*) and unspliced (*u*) graphs:

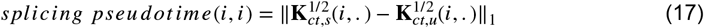

### Software and Datasets

The data and accompanying Mathematica notebooks (Wolfram Mathematica, v13.x for MAC OS X ARM (64-bit), run on 16 GB RAM/M1 or 32 GB RAM/M2 machines) are available at https://github.com/houston-lab/paper-code-files/tree/main. An alternate workflow using a Python Jupyter notebook is also included, modified from a Palantir tutorial using Scanpy v1.9.3/Palantir v1.3.1. ([9]; https://github.com/dpeerlab/Palantir). The original datasets were obtained from online sources: https://github.com/KrishnaswamyLab/MAGIC; https://github.com/dpeerlab/Palantir; https://cellrank.readthedocs.io/en/stable/api/datasets.html. SPRING (https://github.com/AllonKleinLab/SPRING) was run in a Python 2 environment without additional installations. Code for exporting to SPRING is found in the example notebooks.

## Acknowledgements

The author wishes to thank the organizers and other participants of the 2018 Jamboree for *Xenopus* Embryonic Cell Types at the Janelia Research Campus of the Howard Hughes Medical Institute [51] for inspiring an interest in single-cell embryology.

