## Supplemental Figures for "Alternative approaches to single-cell trajectory inference using a commute time matrix"

### Supplemental Figure Legends

**Fig. S1. Quantification of gene interactions using DREMI with commute time imputation.** (A) Density plots of conditional probability for gene expression conditional on expression of *VIM* during EMT. *SDC1* decreases during EMT and has negative correlation ($\rho$) but high DREMI score. *ROBO4* is low during EMT transition but high early and late and thus has no correlation with *VIM* expression. Mutual Information (MI) and DREMI scores are high however. *ZEB-VIM* interactions are shown before and after commute time imputation. (B) Additional analyses of genes predicted to be regulated by ZEB1 [26].

**Fig. S2. Commute time imputation robustness to drop-out and downsampling.** (A) Scatterplots showing imputed data of *VIM-FN1* expression (left), expression after randomly setting 95% of matrix values to zero (middle), re-imputation of the dropout values using the commute time kernel. (B) Scatterplots of imputed *VIM-MYC* expression (left) and the values following four-fold downsampling from 7523 cells to 1881 cells.

**Fig. S3. Commute time imputation and pseudotime applied to human bone marrow differentiation.** (A) Comparison of primary (left) and secondary (right) $k$-NN graphs showing *GATA1* expression. (B) Representation of gene interactions in hematopoietic cell fate choices using commute time imputed expression. Left, erythroid and megakaryocytic markers, highlighted by megakaryocytic marker *ITGA2B;* Middle, dendritic and monocyte markers, highlighted by monocyte marker *MPO;* erythroid and myeloid markers, highlighted by stem cell marker *CD34*. (C) Pseudotime comparison using DPT, Palantir and commute time, using the same root cell in the hematopoietic stem cell population, plotted on the secondary $k$-NN graph. (D) Scatterplot showing correlation of Palantir and commute time pseudotime; some divergence for the most distant pseudotime is evident.

**Fig. S4. Pancreatic lineage marker gene expression.** Force-directed graph plots of E15.5 pancreatic cells highlighting cells expressing lineage-specific and progenitor markers. Cell are colored by scaled, commute time-imputed expression; differentiated endocrine lineages are indicated by the corresponding Greek letters. $\beta$, beta, *Ins2;* $\alpha$, alpha, *Gcg*; $\gamma$, gamma, *Ppy/Pyy*; $\delta$, delta, *Sst*; $\epsilon$, epsilon, *Ghrl*; early (*Neurod2*) and late (*Fev*) endocrine progenitors; ductal cells (*Sox9*). The upper left panel shows the plot of the primary graph, the remainder show the commute time embedding secondary graph.

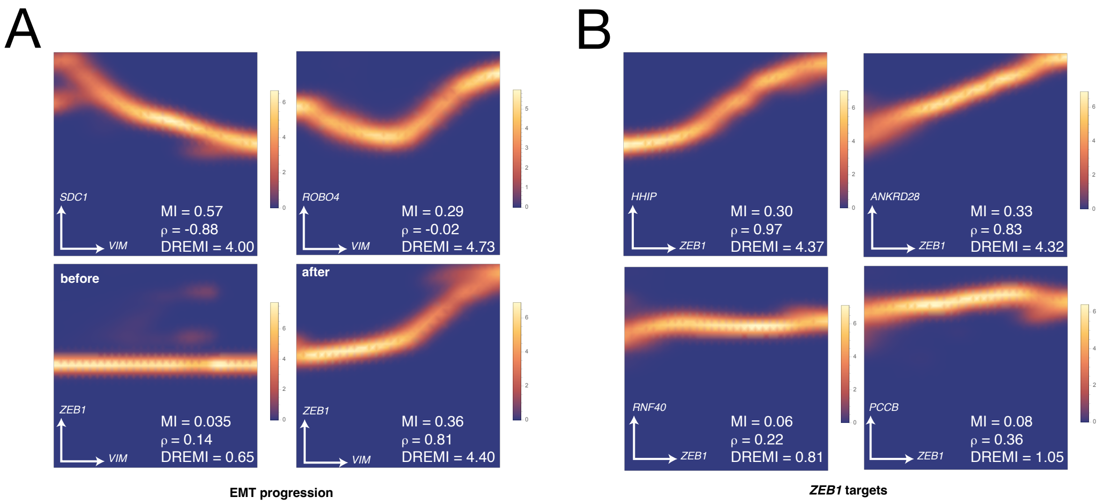


Houston Fig. S1

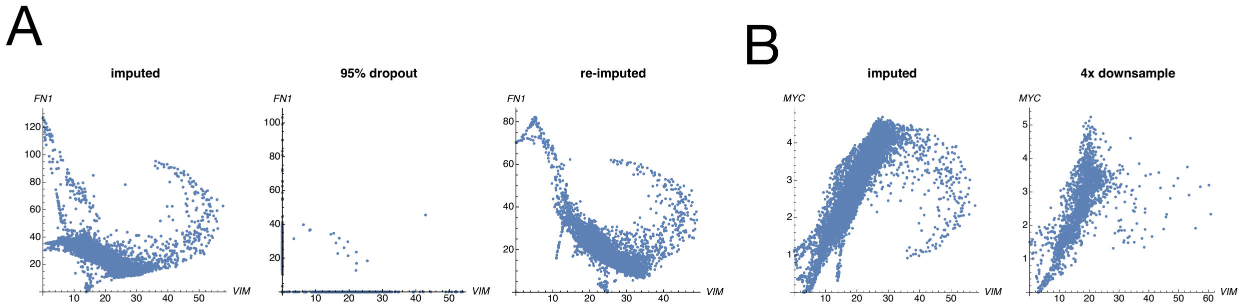


Houston Fig. S2

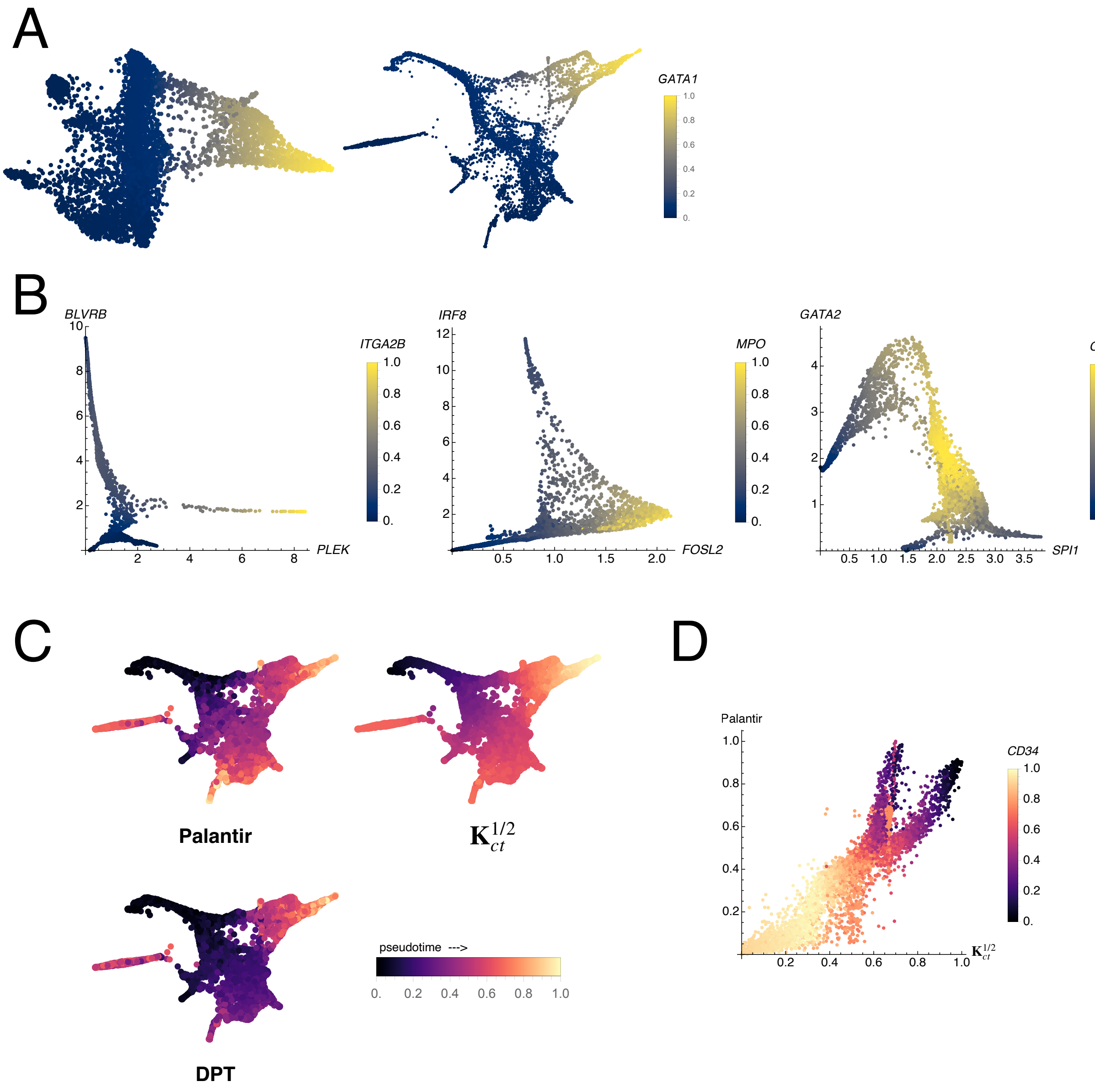


Houston Fig. S3

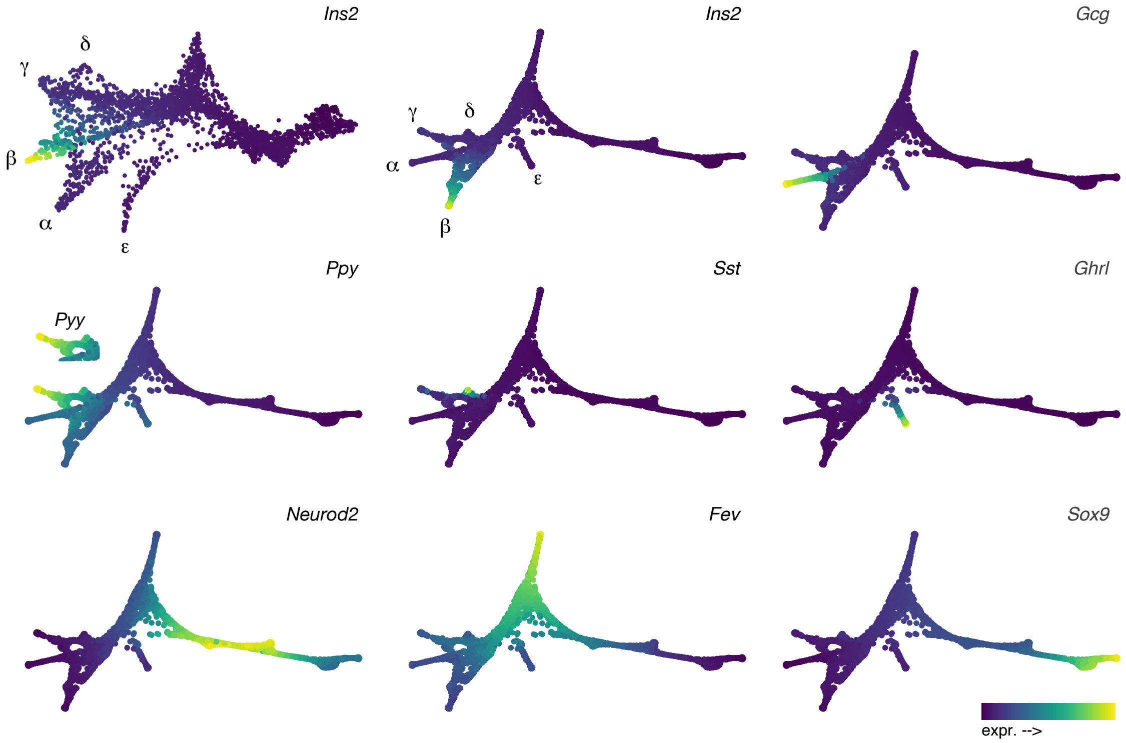


Houston Fig. S4
